## Supplementary Figure for "Assessing the co-variability of DNA methylation across peripheral cells and tissues: implications for the interpretation of findings in epigenetic epidemiology"

**Supplementary Figure 1.** Heatmap of the first ten DNA methylation principal components across five purified blood cell types and three peripheral tissues (whole blood, buccal epithelial cells and nasal epithelial cells). Shown is the mean principal component value for samples grouped by cell- or tissue-type. Each row represents a principal component, with the percentage of variance explained in brackets. Each column represents a sample cell or tissue type.

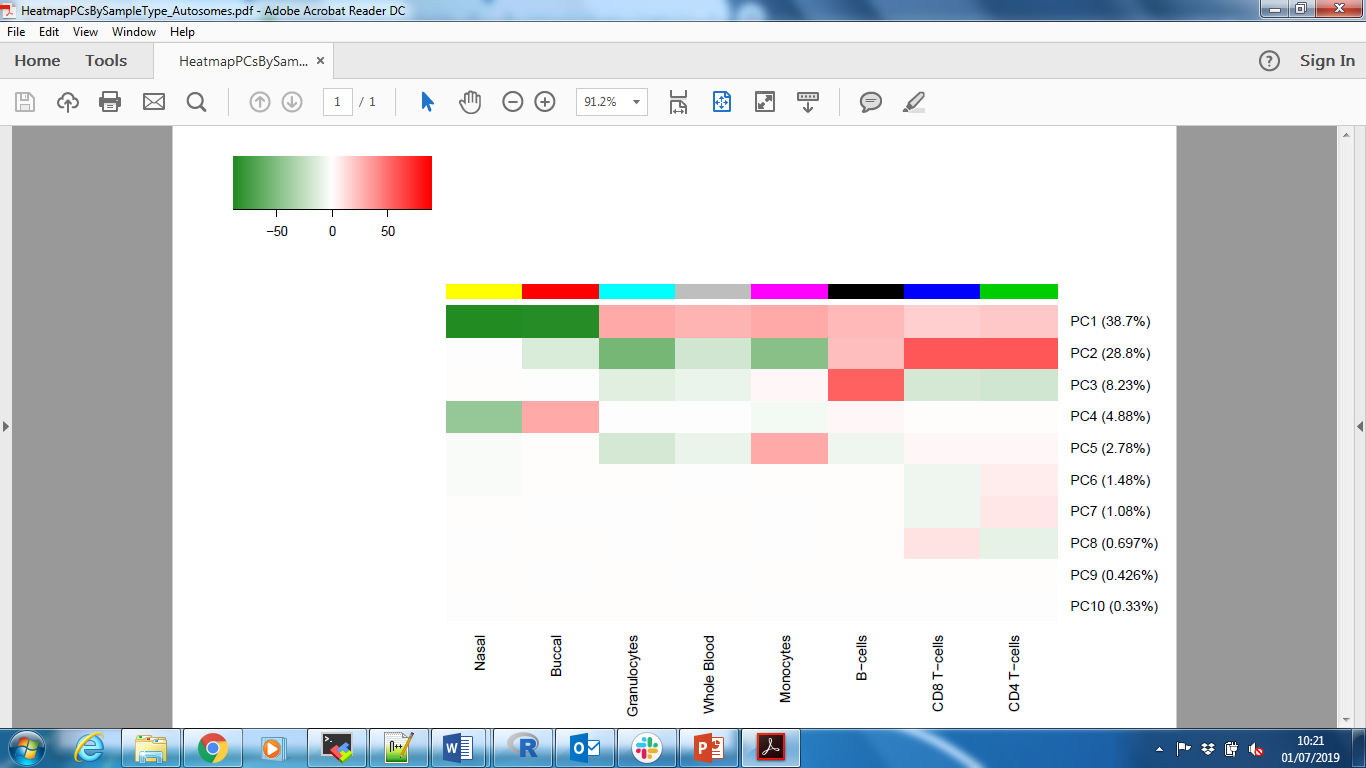

**Supplementary Figure 2.** Density plot of DNA methylation levels across the 784,726 autosomal DNAm sites included in our analysis, separated by tissue and cell-type. Shown is the mean level of DNAm at each site across all individuals.

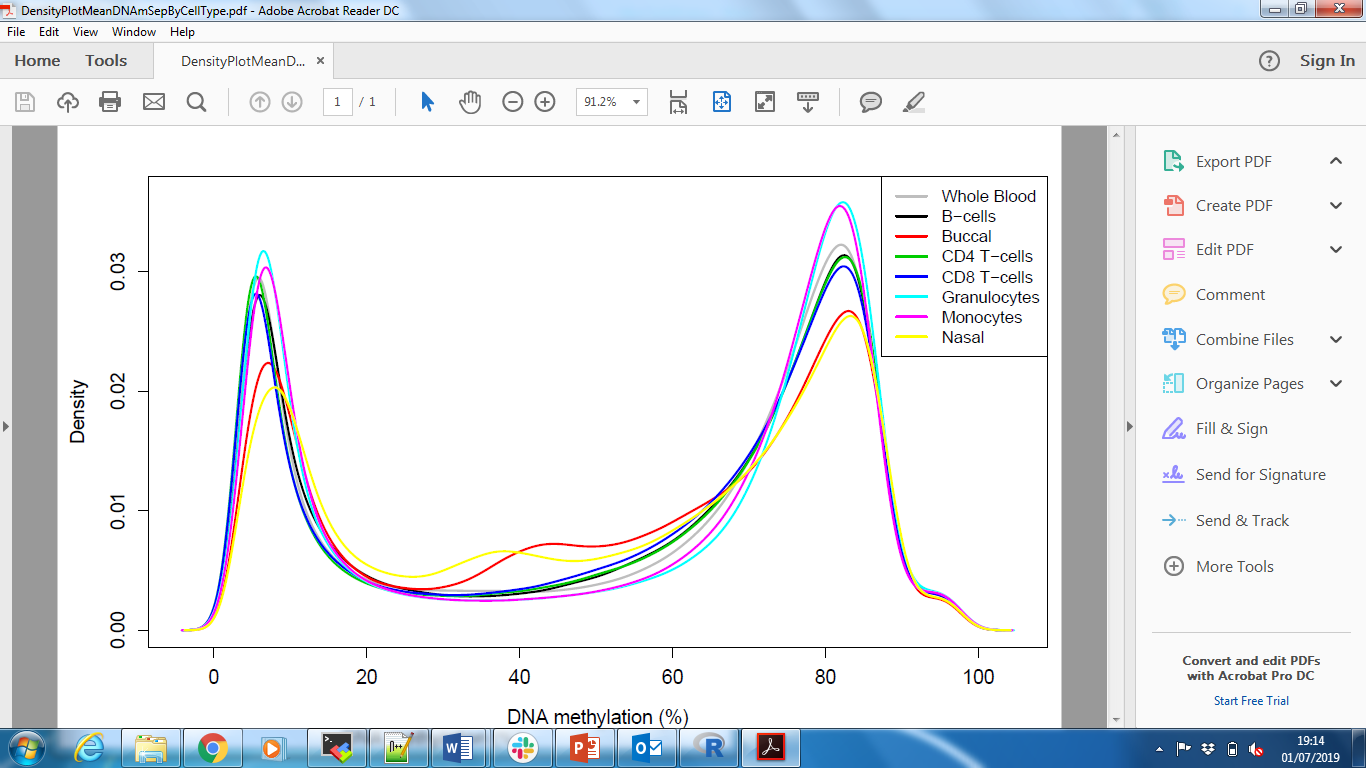

**Supplementary Figure 3.** Barplot showing the proportion of differentially methylated positions (DMPs) shared between tissue and cell-types. For each tissue the sites identified as differentially methylated relative to whole blood were categorized into those that are uniquely different in that sample type or shared with at least one other cell- / tissue-type. Unique DMPs were defined as those where the t-statistic comparing each sample type to whole blood were significant (P < 0.05) for only a single tissue. These two barplots present the A) number and B) percentage of unique and shared DMPs.

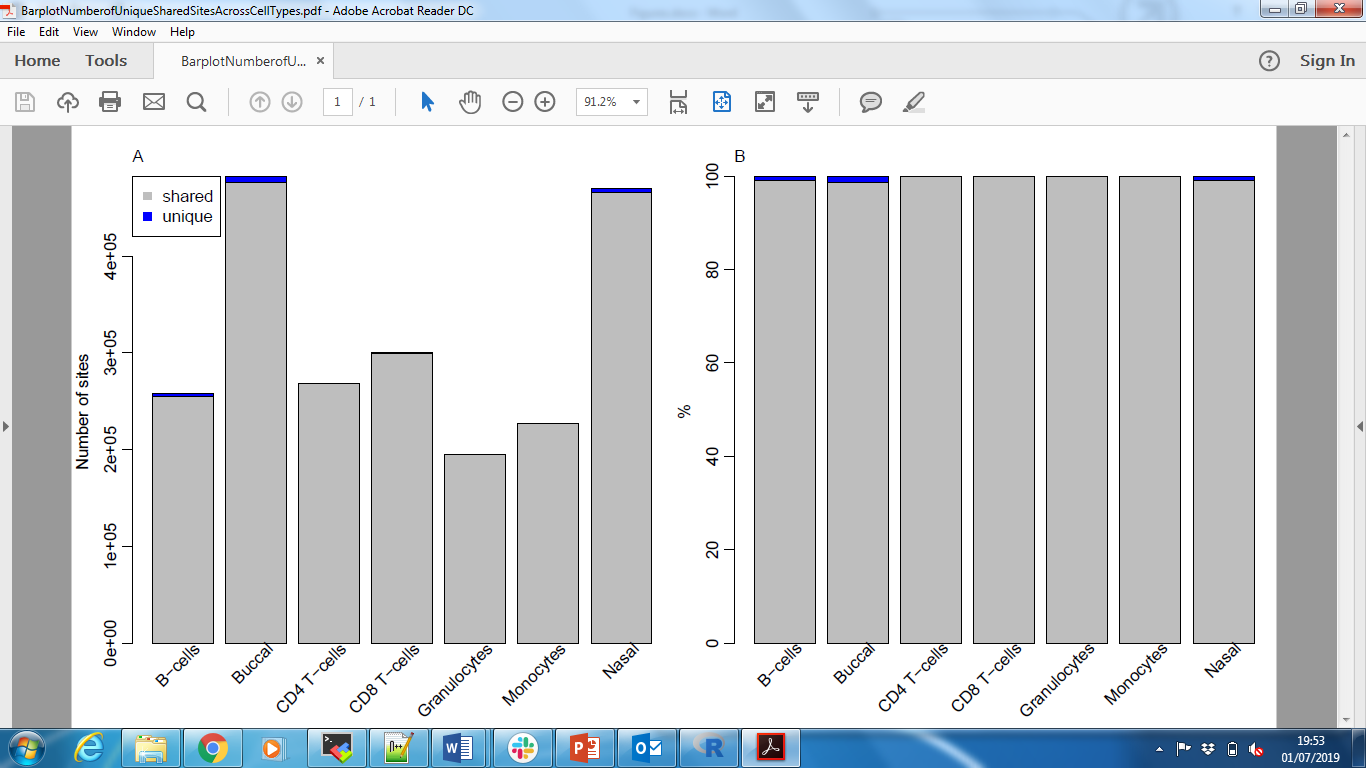

**Supplementary Figure 4. Histogram of the number of cell- and tissue-types in which each DMP is different relative to whole blood.** Taking all sites identified as having a significantly different level of DNA methylation in at least one cell or tissue types (n = 594,056; ANOVA P < 9x10^-8^), we considered t-statistics comparing each sample type and whole blood to count the number of cell/tissue types each site was significantly different in (P < 0.05).

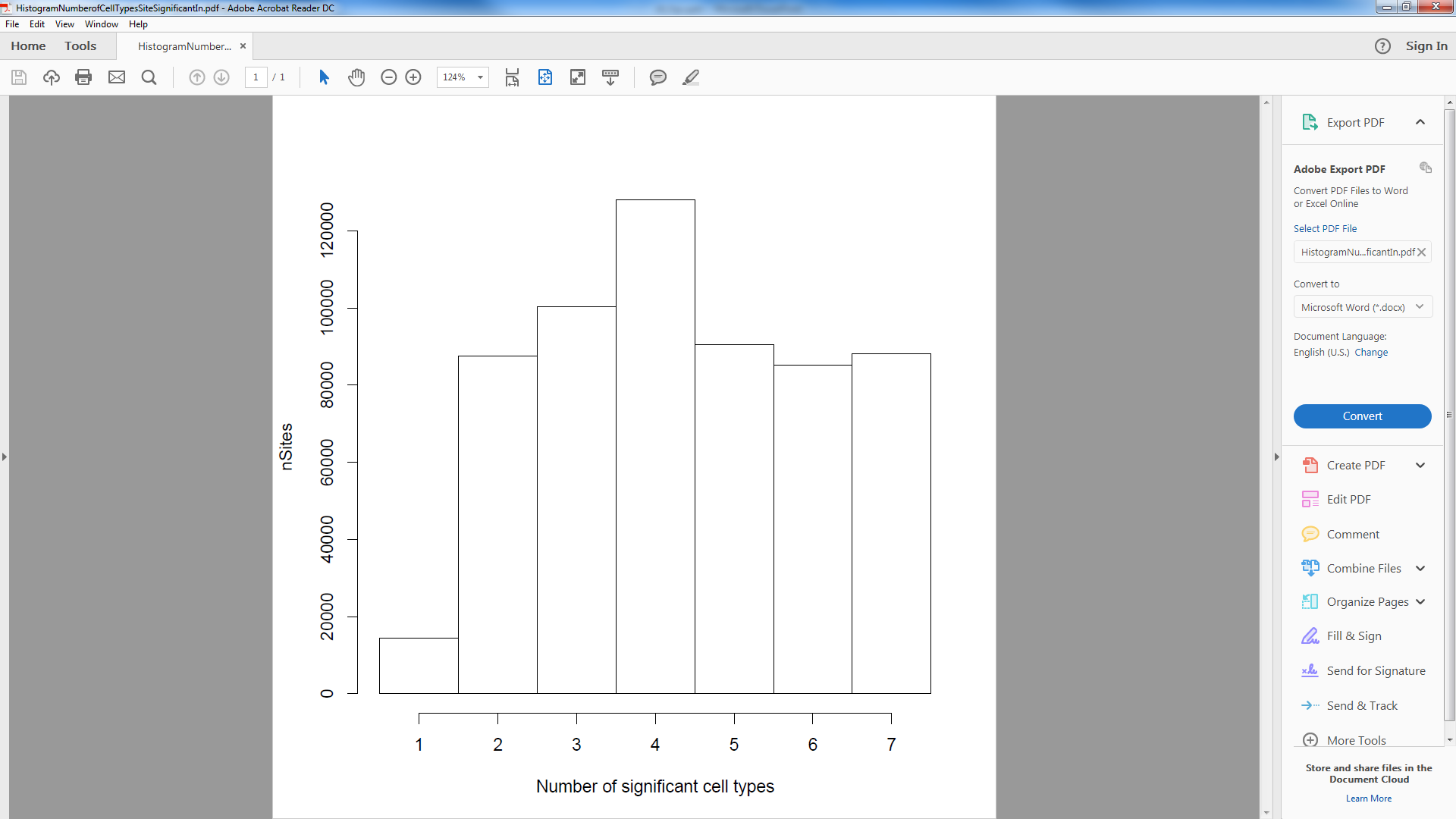

**Supplementary Figure 5. Heatmap showing the overlap between cell- / tissue-types of sites differentially methylated between sample types.** Taking all sites identified as having a significantly different level of DNA methylation in at least one cell or tissue type (n = 594,056; ANOVA P < 9x10^-8^) we considered t-statistics comparing each sample type and whole blood to determine which cell/tissue types each site was significantly different in (P < 0.05). Each box in this heatmap represents the percentage of significantly different DNAm sites that are shared between two cell/tissue types.

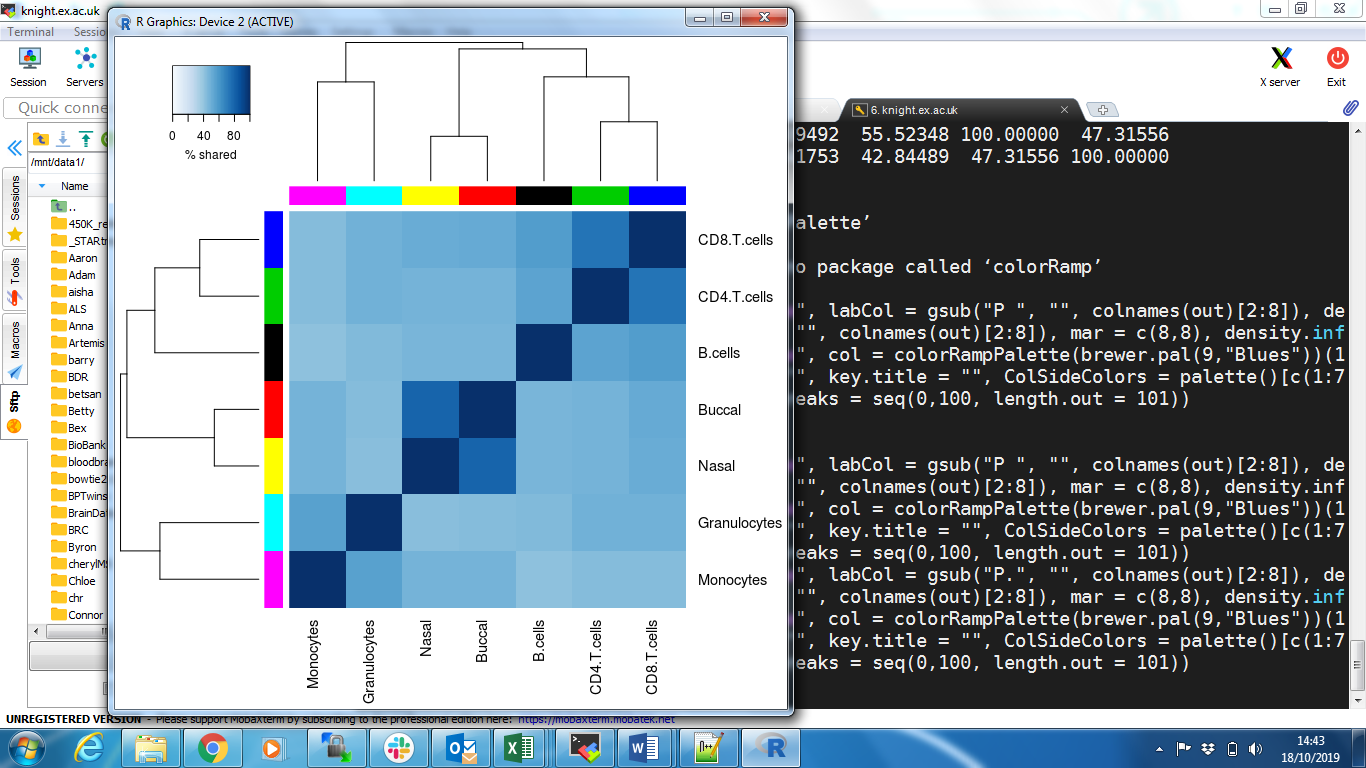

**Supplementary Figure 6: Density plot of the variation in DNAm for each cell- / tissue-type**. Shown across all autosomal DNAm sites included in our analysis is the distribution of the standard deviation at each site. Each sample-type is represented by a different coloured line. This figure shows that in general across the EPIC array, DNA methylation measured in buccal or nasal samples is more variable across individuals than DNA methylation measured in whole blood and individual blood cell types.

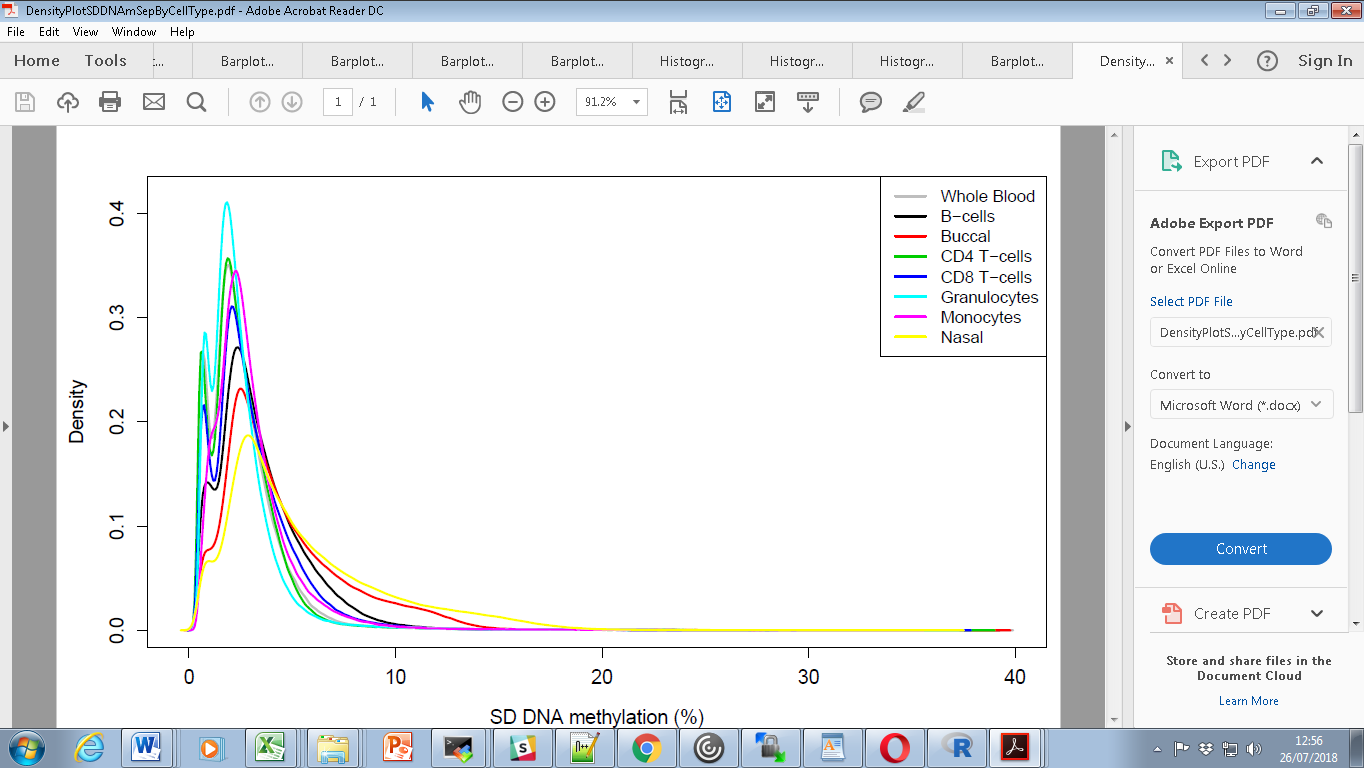

**Supplementary Figure 7**: Scatterplot comparing the variance (standard deviation) in DNA methylation between tissues and cell-types. Shown is data for all autosomal DNAm sites on the EPIC array. Above each plot is the Pearson correlation coefficient.

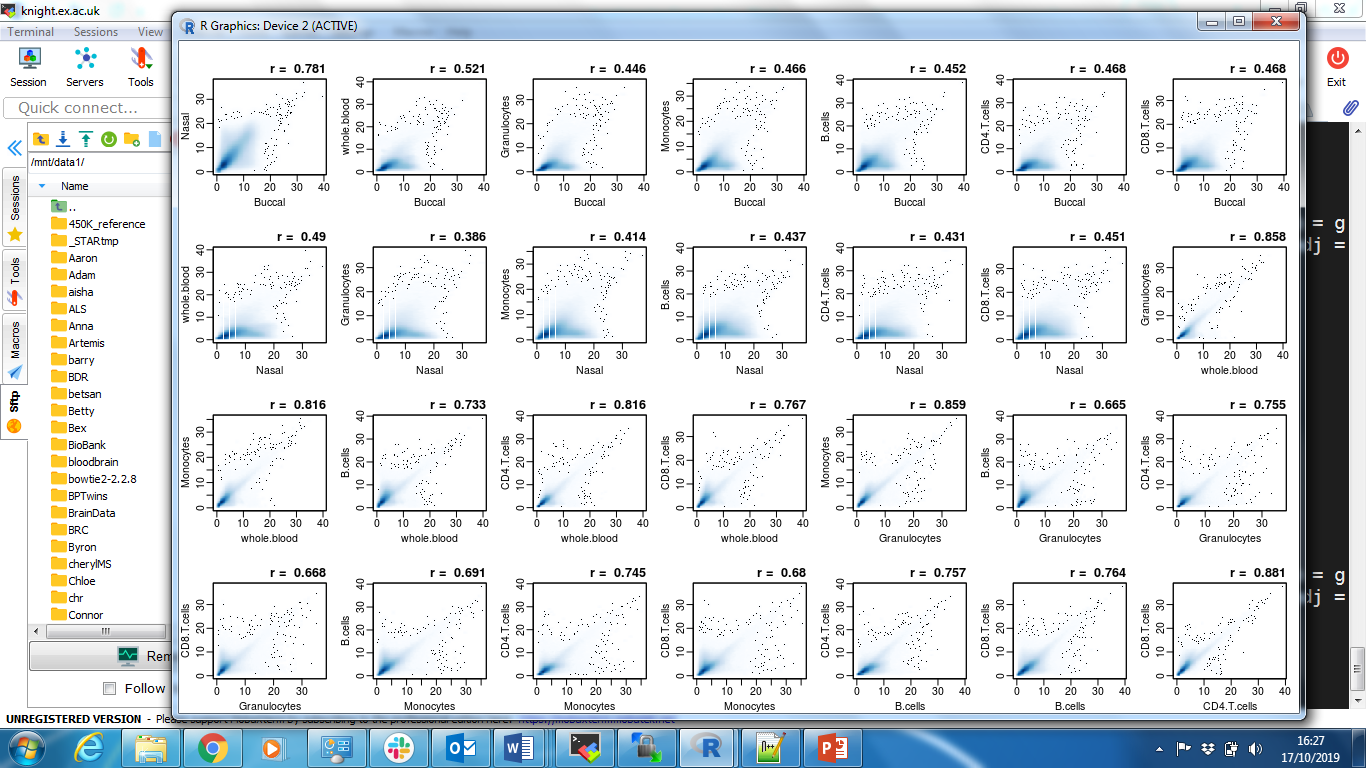

**Supplementary Figure 8**: Density plot of the variation in DNAm for each tissue or cell-type for sites that are variable between sample types. Each sample-type is represented by a different coloured line. This plot shows that sites with significant variance across tissues and cell types are driven by increased variance in buccal and nasal samples compared to whole blood and blood cell types.

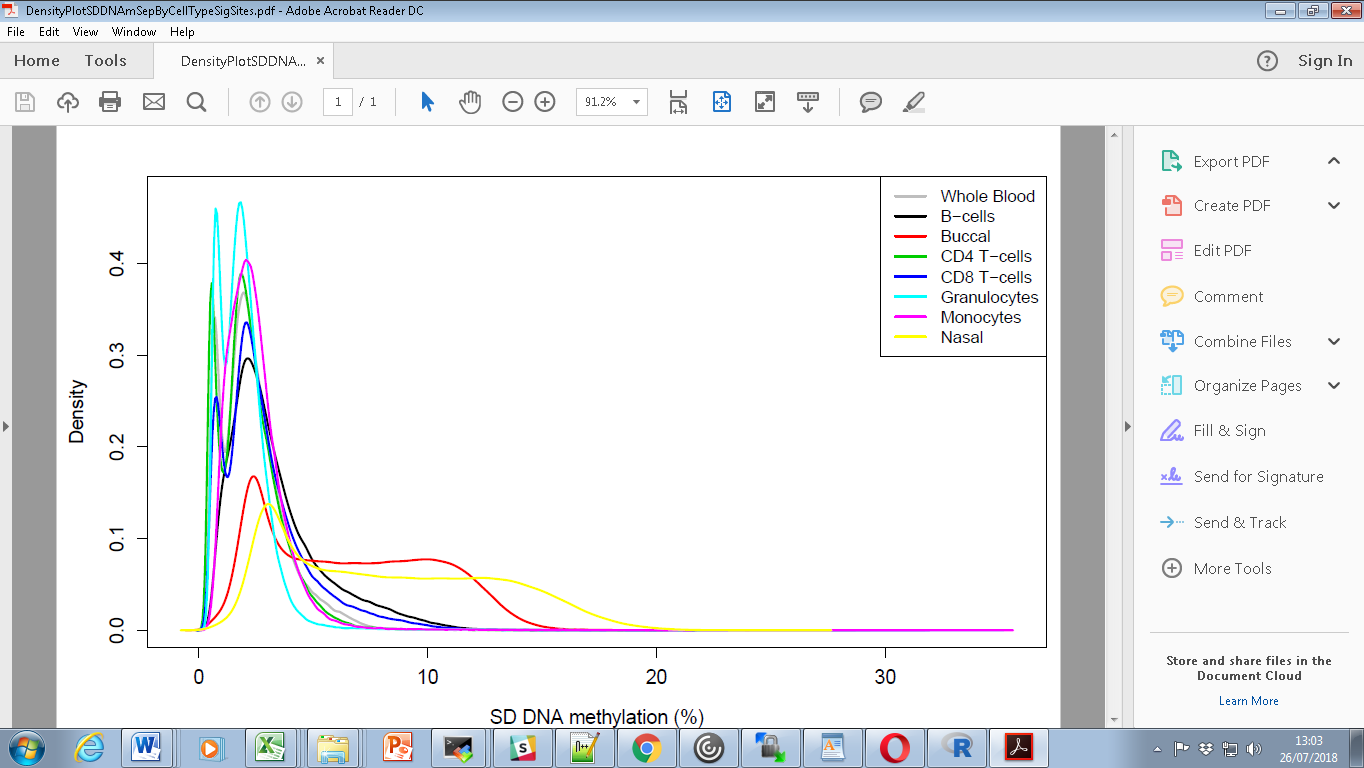

**Supplementary Figure 9. Scatterplot of the variance in DNA methylation (SD) between all pairs of tissue and cells across DNAm sites with significantly difference levels of variation (**n = 196,104). Above each plot is the Pearson correlation coefficient.

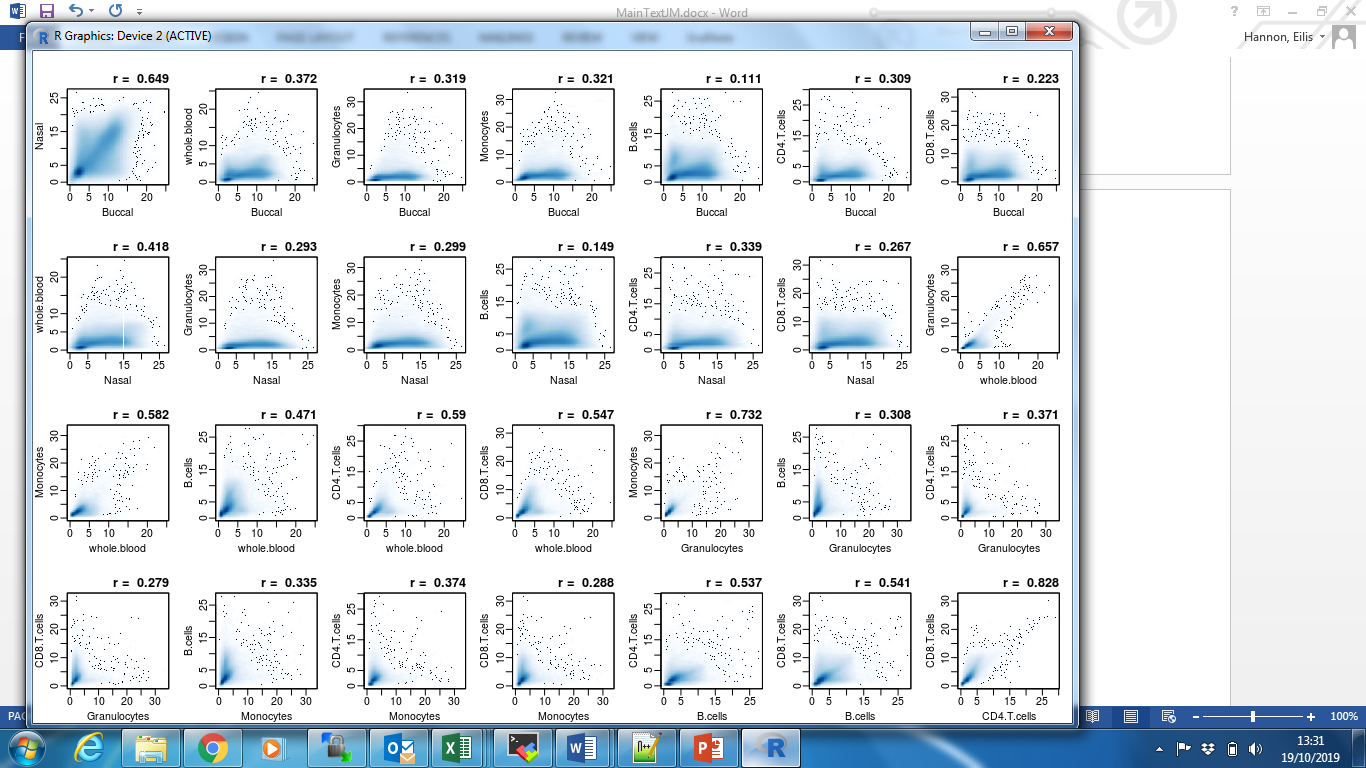

**Supplementary Figure 10. Heatmap of the variance in DNA methylation (SD) between all pairs of tissue and cell-types for DNAm sites shown to vary across sample types (**n = 196,104).

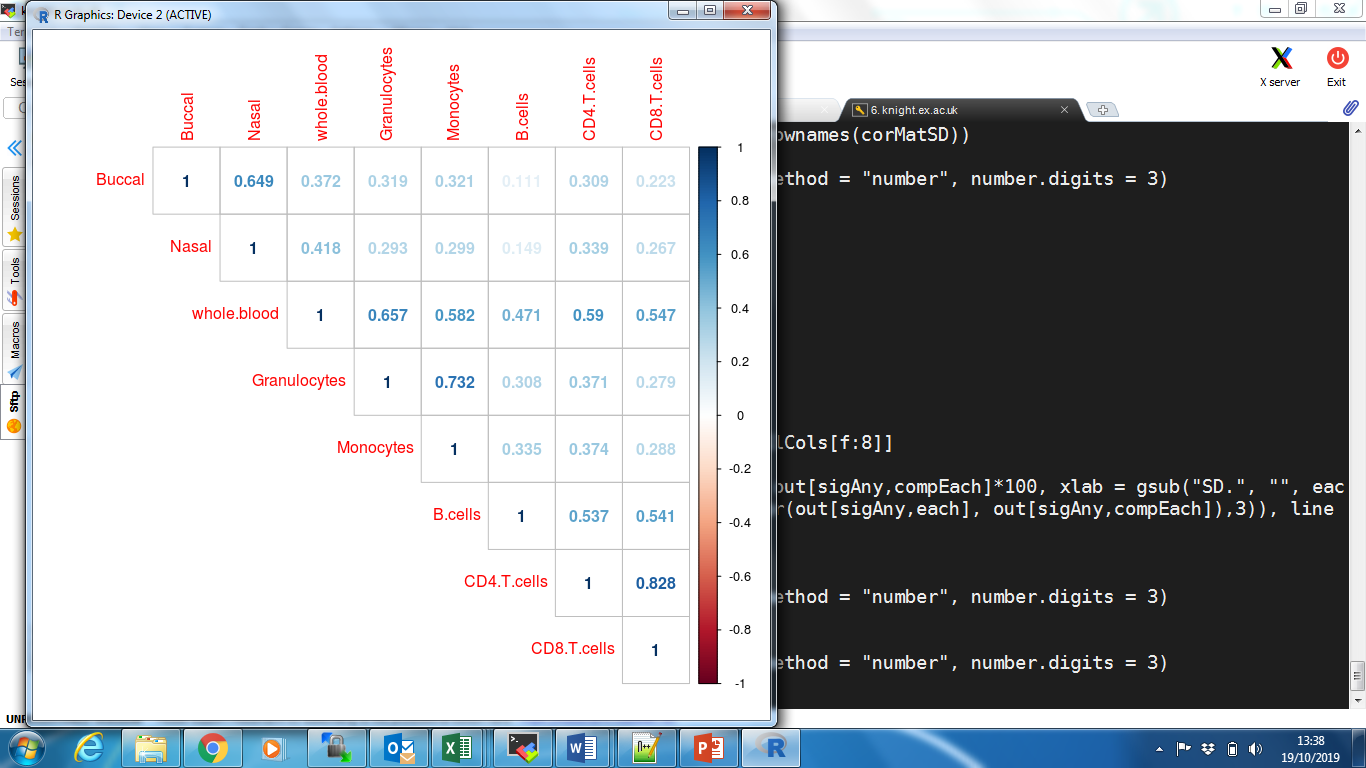

**Supplementary Figure 11: Inter-individual variation in DNA methylation in whole blood is correlated with variation in isolated blood cell types.** Histograms showing the distribution of correlation coefficients between DNA methylation in whole blood and the five blood cell types (A) B-cells; B) CD4 T-cells; C) CD8 T-cells; D) monocytes and E) Granulocytes. The vertical blue dashed line indicates a correlation coefficient of 0. For all five cell types the distribution of correlation coefficients is skewed to the right.

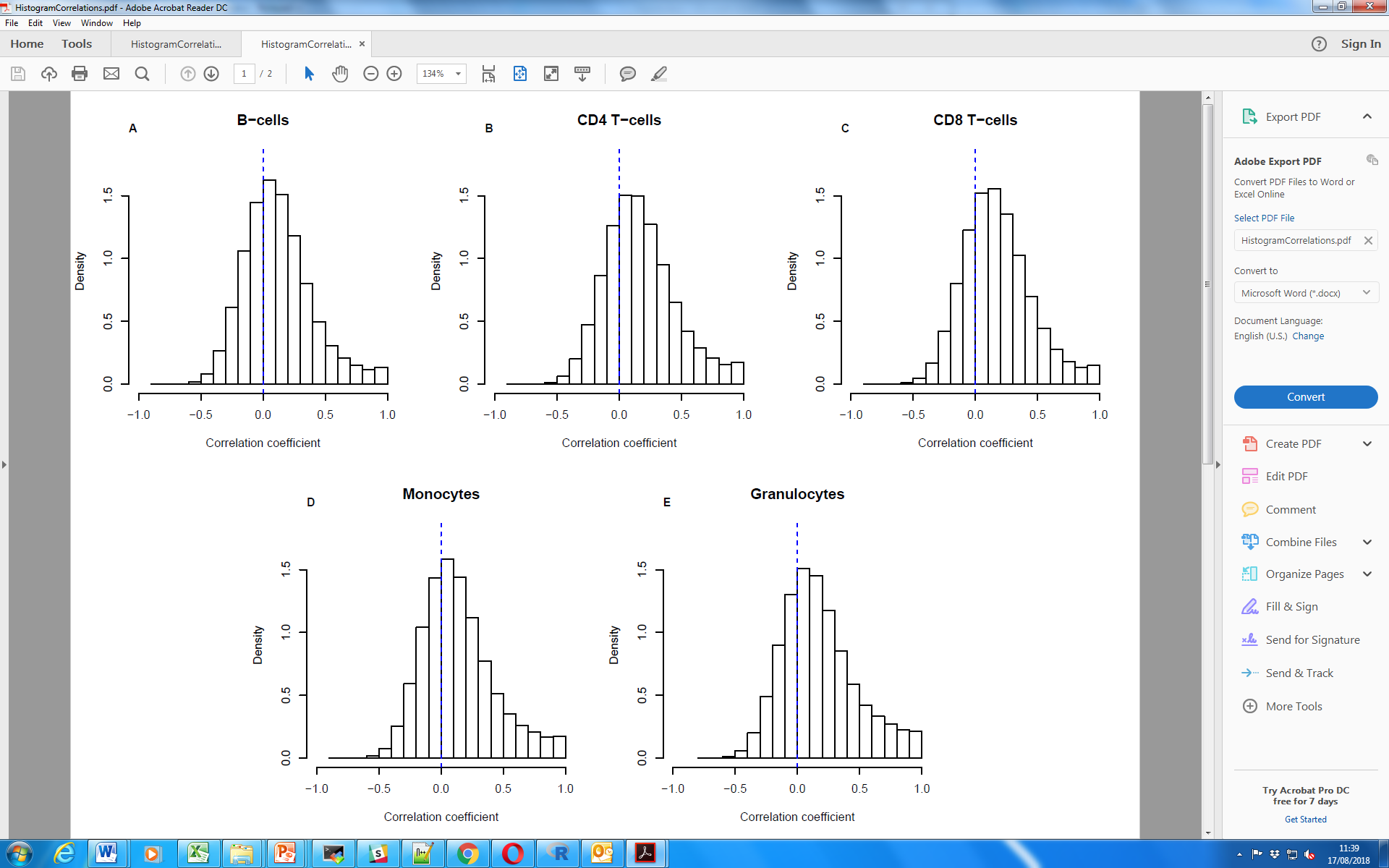

**Supplementary Figure 12. Histogram showing the number of individual blood cell types that explain at least 20% of the variance in whole blood.**

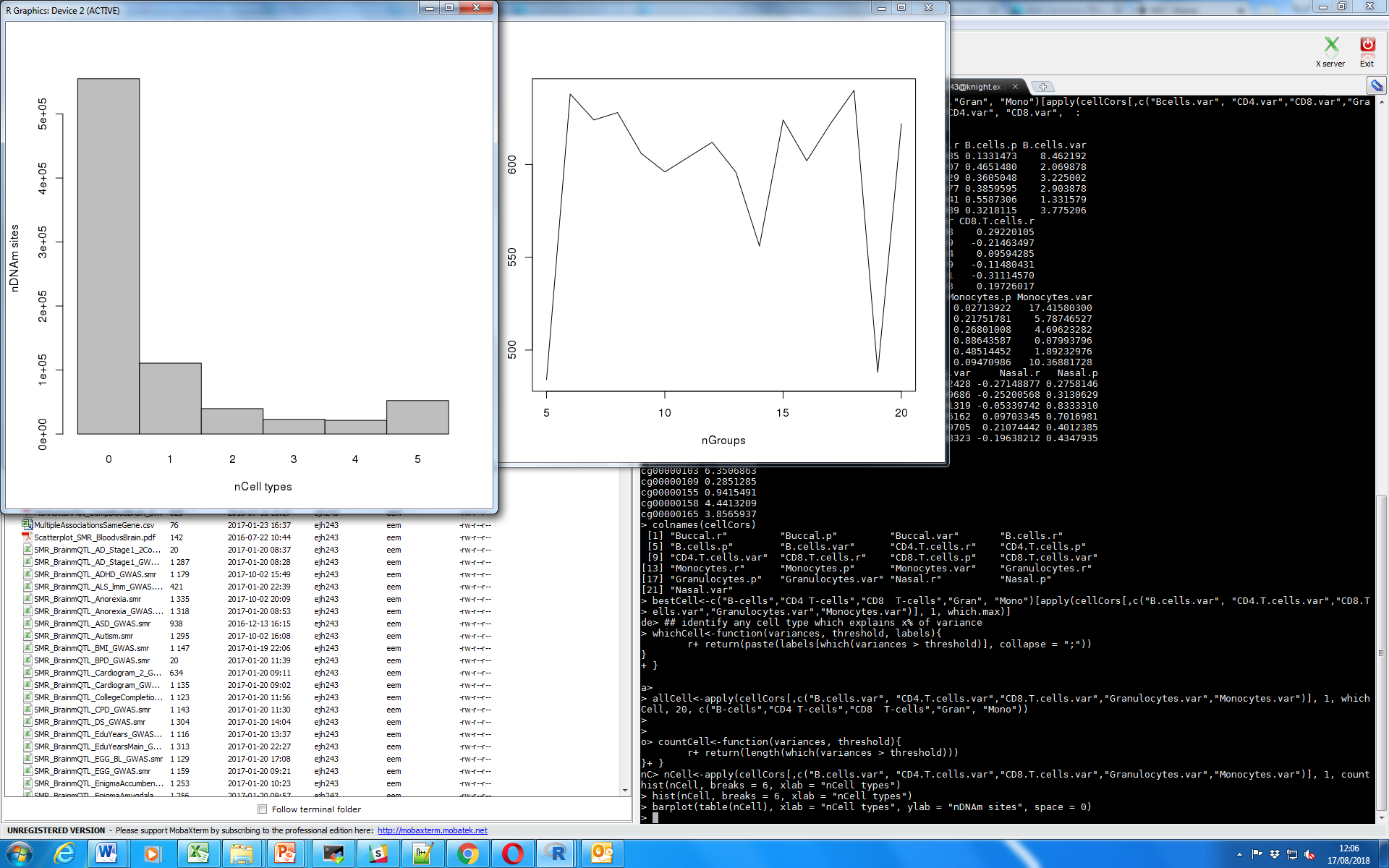

**Supplementary Figure 13. Boxplot of variance explained in whole blood for each cell type separately where DNAm sites are split by mean DNA methylation level and variability.**

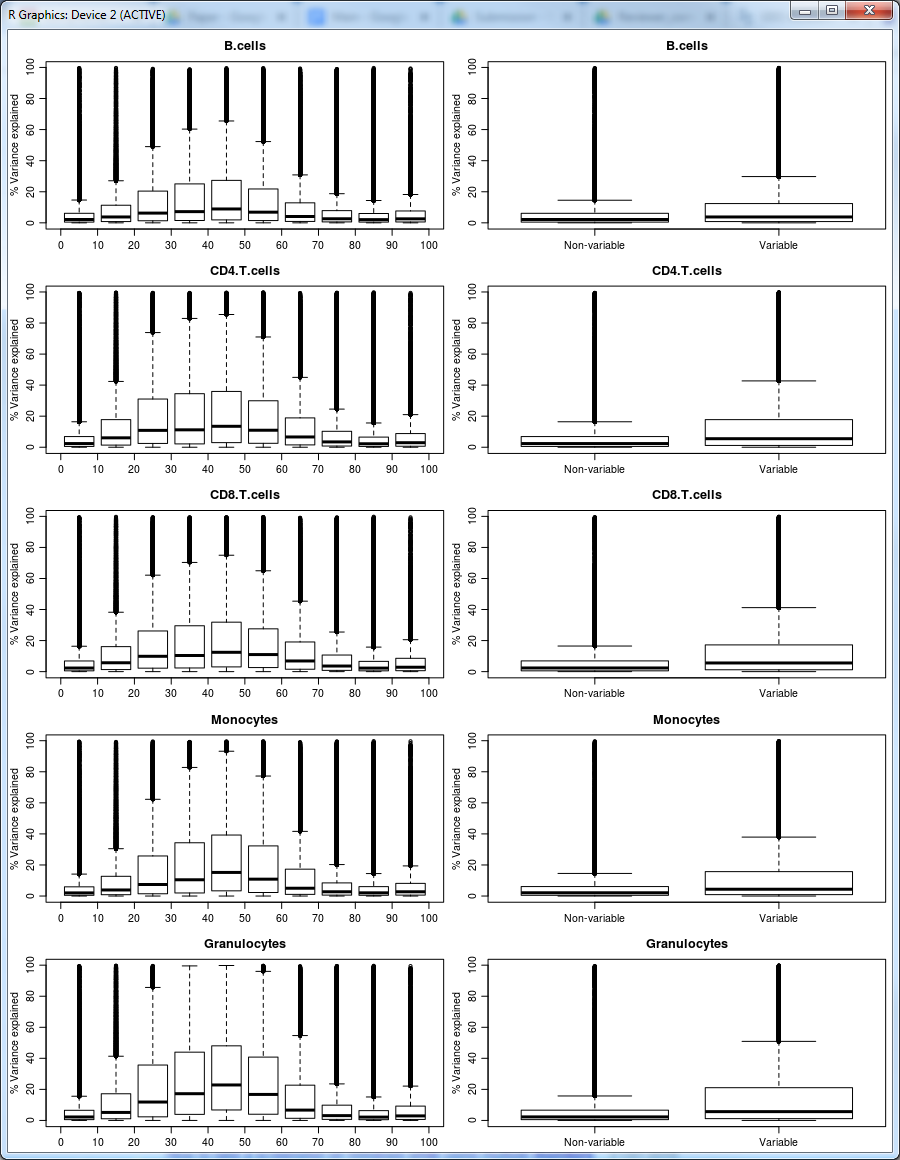

**Supplementary Figure 14: Inter-individual variation in different blood cell types predicts inter-individual variation in whole blood at the same sites.** Scatterplots comparing blood-cell type correlations between cell types. The colour of the point indicates the density of observations at that position ranging from gray (low) to yellow (high).

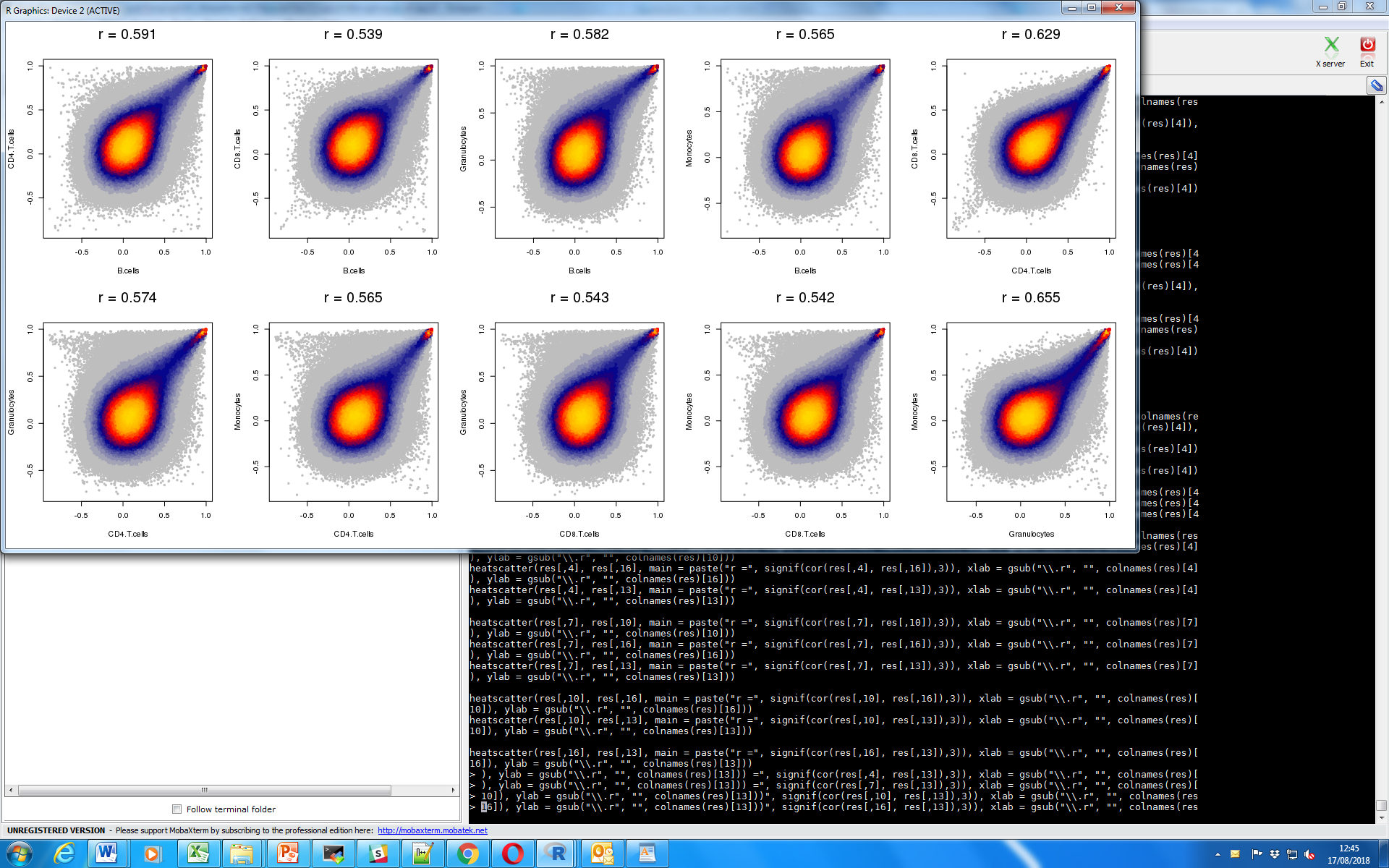

**Supplementary Figure 15. Histogram of variance explained in whole blood by all five cell types together.**

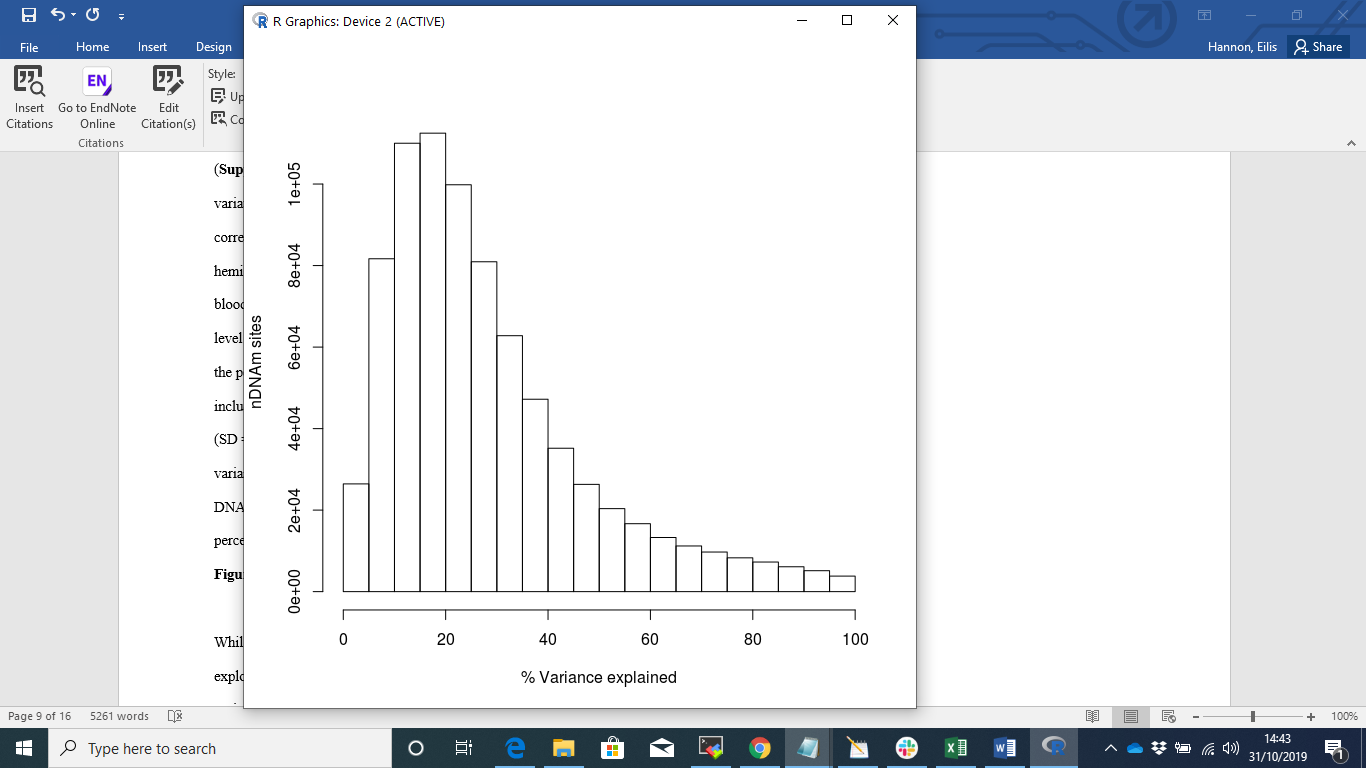

**Supplementary Figure 16: Boxplots of variance explained in whole blood by all five cell types together split by A) mean DNA methylation level and B) variability.**

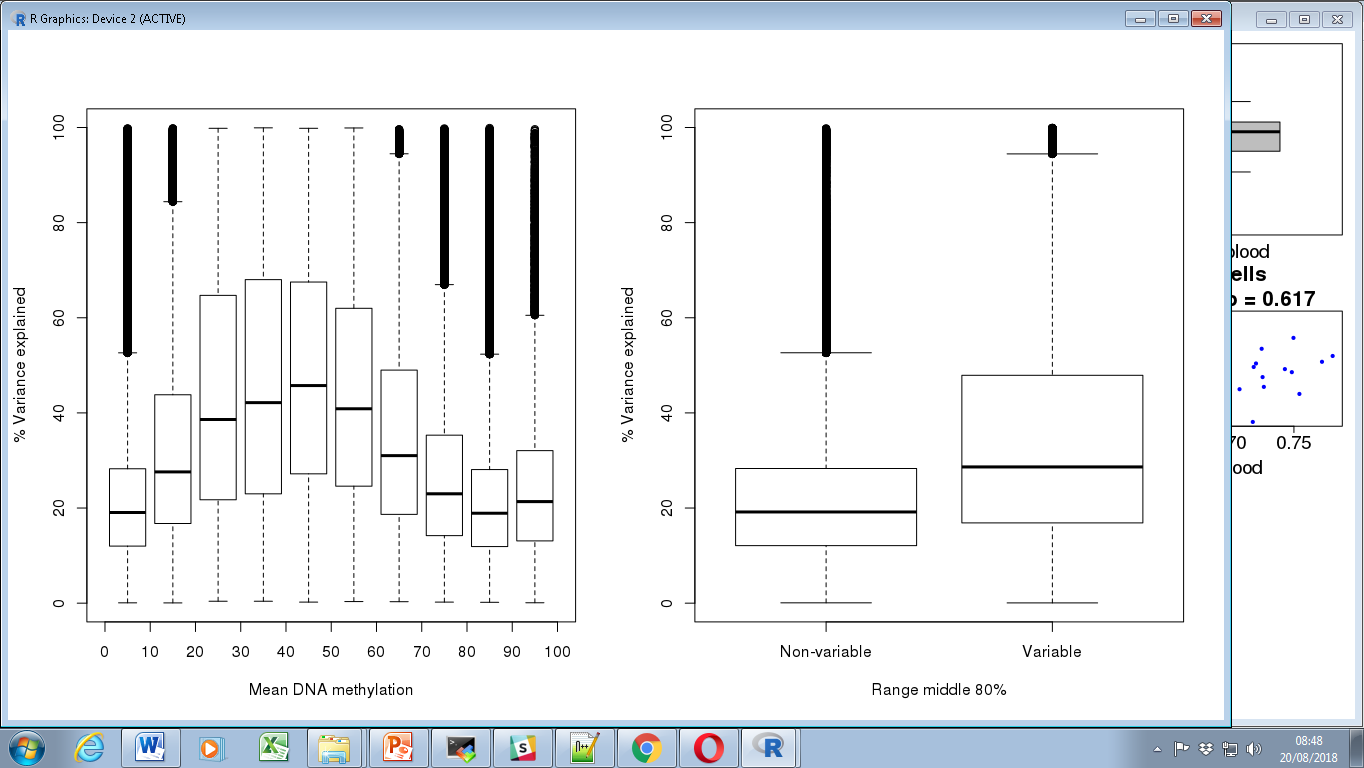
